# Biofilm formation in *Zymoseptoria tritici*

**DOI:** 10.1101/2023.07.26.550639

**Authors:** Tegan E. Tyzack, Christian Hacker, Graham Thomas, Helen N. Fones

## Abstract

*Zymoseptoria tritici* is an economically damaging fungal pathogen of wheat which is able to survive for long periods on the leaf surface. In this environment, the fungus is exposed to many stresses, including fluctuations in temperature, UV radiation, drying, and foliar fungicide applications. We observed biofilm-like cell aggregations on the surface of wheat leaves infected with *Z. tritici* in both compatible and non-compatible interactions. A literature search revealed few examples of plant pathogenic fungi forming biofilms, but demonstrated that such biofilms have a number of key characteristics, which are shared with other biofilm-forming microbes. These include adhesion of cells to the substrate, production of extracellular matrix, altered gene expression and increased tolerance to various stresses. We used a combination of microscopy, qRT-PCR and stress tolerance assays to determine whether putative *Z. tritici* biofilms possessed these diagnostic characteristics. We show that *Z. tritici* biofilms resemble in form and function those formed by other filamentous and dimorphic fungi, producing extra-cellular matrix in which cells become embedded, creating layers of hyphal bundles and blastospores. These biofilms show increased tolerance to drying and high temperature. There is also isolate-dependent resistance to reactive oxygen stress and the fungicide carboxin. Taken together, these findings demonstrate that *Z. tritici* is indeed able to produce genuine biofilms.

## Introduction

Biofilms are most simply described as layers of microbial cells growing on a substrate. The existence of such films was first noted in bacteria, partly as a result of the observation that available surface area positively affects bacterial population growth in samples of seawater[1], while the term ‘biofilm’ was coined later[2] to describe the coating of bacteria found on rocks in freshwater. More recently, the study of bacterial biofilms has gathered pace due to their clinical relevance; bacterial biofilms are common features of disease and frequently colonise indwelling medical devices[3]. Biofilm cells show increased resistance to antibiotics and can make infections harder to eradicate[4–6]. The ability to form biofilms is ubiquitous in bacteria and occurs in a wide range of environments[7]. Advancements in our understanding have uncovered the key features of these microbial structures. In biofilms, cells are attached to their substrate and to each other and are embedded in an extracellular matrix (ECM) consisting of extracellular polysaccharides such as alginate, proteins, extracellular DNA and lipids[8, 9]. This ECM may be architecturally complex, with channels allowing diffusion of water, oxygen and nutrients[8, 10]. Despite this, biofilms often include cells with reduced access to external nutrients and thus reduced metabolic activity. This may extend to dormancy or even cell death[11]. Further, biofilm formation is associated with increased antimicrobial resistance (AMR)[12, 13].

Fungi can form structures analogous to the biofilms seen in bacteria. This is particularly clear in the case of yeasts[14, 15]. Again, human pathogens have received the most research attention, with biofilm formation in *Candida* species, which also colonise surfaces of indwelling medical devices, particularly well understood [15–18]. *Candida* biofilms are formed from a mixture of hyphal and yeast-like cells; their formation begins with the adhesion of yeast-like cells to a substrate, with a layer of interconnected hyphae developing above that base [18–21]. As with bacterial biofilms, the cells are embedded in an ECM composed of polymeric substances[17] and show increased stress tolerance and resistance to antimicrobials[22]. AMR is a key feature of *Candida* biofilms in clinical settings[19, 21, 23]. The precise mechanism underpinning AMR is unclear. Studies have provided evidence for hypotheses including sequestration of antimicrobials by ECM components[24, 25], higher expression of efflux pumps in biofilm compared to pelagic cells[15, 26] and the effect of lowered metabolic rates inside biofilms on antimicrobial metabolism and efficacy[27, 28]. It seems likeliest that some combination of these factors is responsible, with the most important factor depending on the stage of biofilm development. This can be seen in that an increase in fungicide resistance is seen in *Candida* cells after attachment, but before the development of ECM; during this period, ABC transporter efflux pumps are upregulated and changes in expression of metabolism genes are detected [15]. However, resistance increases further once the biofilm matures. ECM, specific morphology and further differences in gene expression are implicated, but efflux pumps are thought to be important in the AMR of these mature biofilms[15]. The definition of a biofilm is more complex in filamentous fungi, in which attachment to a substrate is typical and growth dynamics markedly different from bacteria or budding yeasts[14]. However, taking into account the key characteristics of both bacterial and yeast biofilms – namely, that cells are embedded in an ECM, show increased stress tolerance and AMR, and altered gene expression when compared to pelagic cells, it is possible to identify filamentous fungal structures which are best described as biofilm [14, 29]. As with other types of biofilm, those formed by filamentous fungi are best studied where they impinge upon human health, including biofilms formed by opportunistic human pathogens in genera such as *Aspergillus* and *Fusarium*[30].

In *Aspergillus fumigatus*, ECM is highly heterogeneous and predominantly composed of EPS such as *α*-glucans, galactosaminogalactan (GAG) and galactomannan. Other components include polyols, lipids, DNA and proteins[30]. Interestingly, GAG appears to function in many of the key biofilm characteristics, being involved in adhesion to surfaces, biofilm architecture and stability, AFR and host immune evasion[31]. In *Fusarium solani* biofilms, ECM has a similar composition, being primarily composed of extracellular polysaccharides (EPS) with DNA and proteins also present[32]. Importantly, hyphae within the biofilms of filamentous fungi are interlinked, often *via* anastomoses, and interwoven so that, along with the ECM, they form complex architectures complete with channels for the diffusion of oxygen, water and nutrients[29, 32]. In common with other biofilms, the central areas may nevertheless be hypoxic, with dormant and dead cells acting as part of the overall biofilm structure. These help, along with the ECM, to protect live, metabolically active cells from stressors such as fungicides, UV and temperature[31–34]. Development of these filamentous biofilm structures can be characterised as a step-wise process beginning with adhesion, progressing through the formation of microcolonies, which may involve the development of hyphal bundles or layers, anastomosis and ECM secretion, and finally the maturation into a complex biofilm structure of multiple hyphal layers embedded in ECM[29, 32, 33]. Importantly, filamentous biofilms may be disrupted by the *Candida* quorum sensing molecule, farnesol, suggesting that cell-cell communication is important within these structures[30, 35]. AFR and other forms of stress resistance are, again, associated with the ECM and with reduced metabolism at the centre of the biofilm, but there is also evidence of differential expression of ergosterol biosynthetic genes[31]. Ergosterol is a membrane lipid with roles in multiple stress responses and is the target of azole fungicides[36–38]. Overall, it is clear that filamentous fungal biofilms are as complex and as adapted for stress tolerance as bacterial and yeast biofilms, and thus are likely to be of fundamental importance in understanding how fungi persist in challenging environments and overcome the stresses involved in host invasion.

As well as their roles as human pathogens, many filamentous or dimorphic fungi are economically and ecologically important plant pathogens[39, 40]. Plant infection requires survival in potentially hostile environments, since plants are exposed to changing weather conditions and, in agriculture, may be heavily treated with antifungals[39–43]. This makes biofilm growth in plant pathogenic fungi of interest, as it may influence survival in the face of these stresses and thus pathogen evolution and the concerning phenomenon of AFR emergence in agricultural settings[41, 44, 45]. However, due to the relatively recent recognition of the possibility of biofilm formation in filamentous fungi, this area is understudied. A few examples of biofilm formation in plant pathogenic fungi have been reported, in several *Fusarium* species as well as plant-pathogenic *Aspergillus, Verticillium* and *Botrytis*[29, 32, 33, 46–53]. We observed biofilm-like growth on the surface of wheat leaves by certain isolates of the fungal wheat pathogen, *Zymoseptoria tritici. Z. tritici*, the causal agent of Septoria tritici blotch in wheat, is the most important fungal pathogen of temperate-grown wheat, costing the UK alone up to £400M in lost wheat yield and fungicide inputs annually[43]. This dimorphic fungal phytopathogen is unusual in that it is able to survive for several days on the leaf surface[54, 55] prior to the leaf penetration and internal colonisation that is essential for disease[56, 57]. Germination of spores on the leaf surface is asynchronous[55] and surface exploration by hyphae is apparently random[54], so that the length of time spent on the leaf surface may extend for many days[54, 55]. Different isolates show different quantities of surface growth even when they have equivalent virulence[58]. Further, some avirulent isolates show the ability to undergo microcycle conidiation on the leaf surface, reproducing by budding to produce significant biomass that is hypothesised to have a role in survival and dispersal to susceptible hosts or in increasing the availability of these isolates as mating partners to virulent isolates in the later, sexual part of *Z. tritici*’s life cycle[59]. Further, *Z. tritici* has been shown to survive for long periods in soil[60]. These facets of the *Z. tritici* life cycle mean that the fungus is exposed to environmental stresses to an unusual degree for a plant pathogenic fungus. When on the leaf surface, the fungus will encounter fluctuating temperatures, drying, and UV, as well as foliar fungicides. Previously, Duncan & Howard (2000) demonstrated that

*Z. tritici* produces an ECM when grown on Teflon surfaces, although they were unable to demonstrate ECM production *in planta*[61]. In this work, we note that the accretions of fungal growth seen in on leaf surfaces inoculated with avirulent (or “NIRP”) isolates[59] resemble biofilm-like growth and we investigate the possibility that *Z. tritici* is able to form biofilms.

## Results

### Biofilm-like growth can be observed in *Z. tritici* growing on both susceptible and resistant wheat hosts

Observation of cytoplasmic GFP tagged isolates of *Z. tritici* on the surface of wheat leaves showed that, in addition to the commonly described process of germination, surface exploration and stomatal penetration by hyphae (Figure 1A), the fungus can produce ‘mats’ consisting of hyphae and blastospores on the leaf surface. These can be observed in avirulent interactions with resistant wheat (isolate T39 on the bread wheat Galaxie; (Figure 1B)), as we reported previously [59]. Hyphae are commonly visible connecting the densest blastospore formations and at the edges of the fungal mat (Figure 1B). Similar blastospore mats can also be observed in virulent interactions with suspectable wheat (isolate IID2-157 on durum wheat; (Figure 1C)). Representative images of infected leaves at 21 dpi, showing the differing degrees of pycnidiation in the three isolate-wheat combinations, are shown in Figure S1. These mats of fungal material are reminiscent of biofilms produced by certain fungi, such as the mixed hyphal and blastosphore biofilms produced by *Candida albicans*[18].

**Figure 1.**
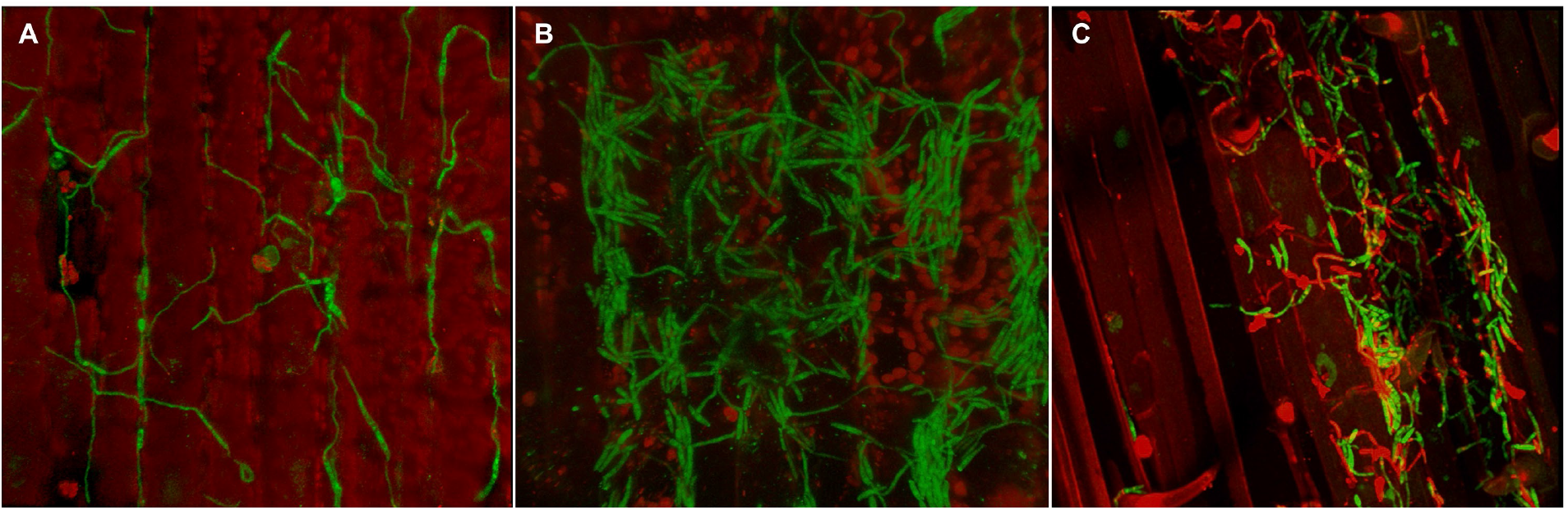
Biofilm-like growth in *Z. tritici*. Representative confocal scanning microscopy images of cytoplasmic GFP-tagged *Z. tritici* isolates growing on wheat hosts. A: Isolate IPO323 on Galaxie bread wheat at 12 dpi. B: Isolate T39 on Galaxie bread wheat at 15 dpi. C: IID2-157 on Volcani durum wheat at 15 dpi, with propidium iodide counterstaining of dead tissues. Image A shows the typical presentation of *Z. tritici* growing over the wheat leaf surface; biofilm-like mats of cells can be observed in images B and C.

Whether these observed fungal structures can be defined as a genuine biofilm depends on the accepted definition of a biofilm in dimorphic plant pathogenic fungi such as *Z. tritici*. We therefore conducted a review of the literature concerning biofilm formation specifically in filamentous or dimorphic fungal plant pathogens, the results of which are summarised in Table 1. It is clear that both hyphae and blastospores can be involved in biofilm formation *in planta* (Table 1). As with biofilms both in other fungi and in bacteria, the defining features include the attachment of cells to a surface, the production of an extracellular matrix surrounding these cells and an increase in resistance to certain stressors when growing as a biofilm (Table 1).

**Table 1.**
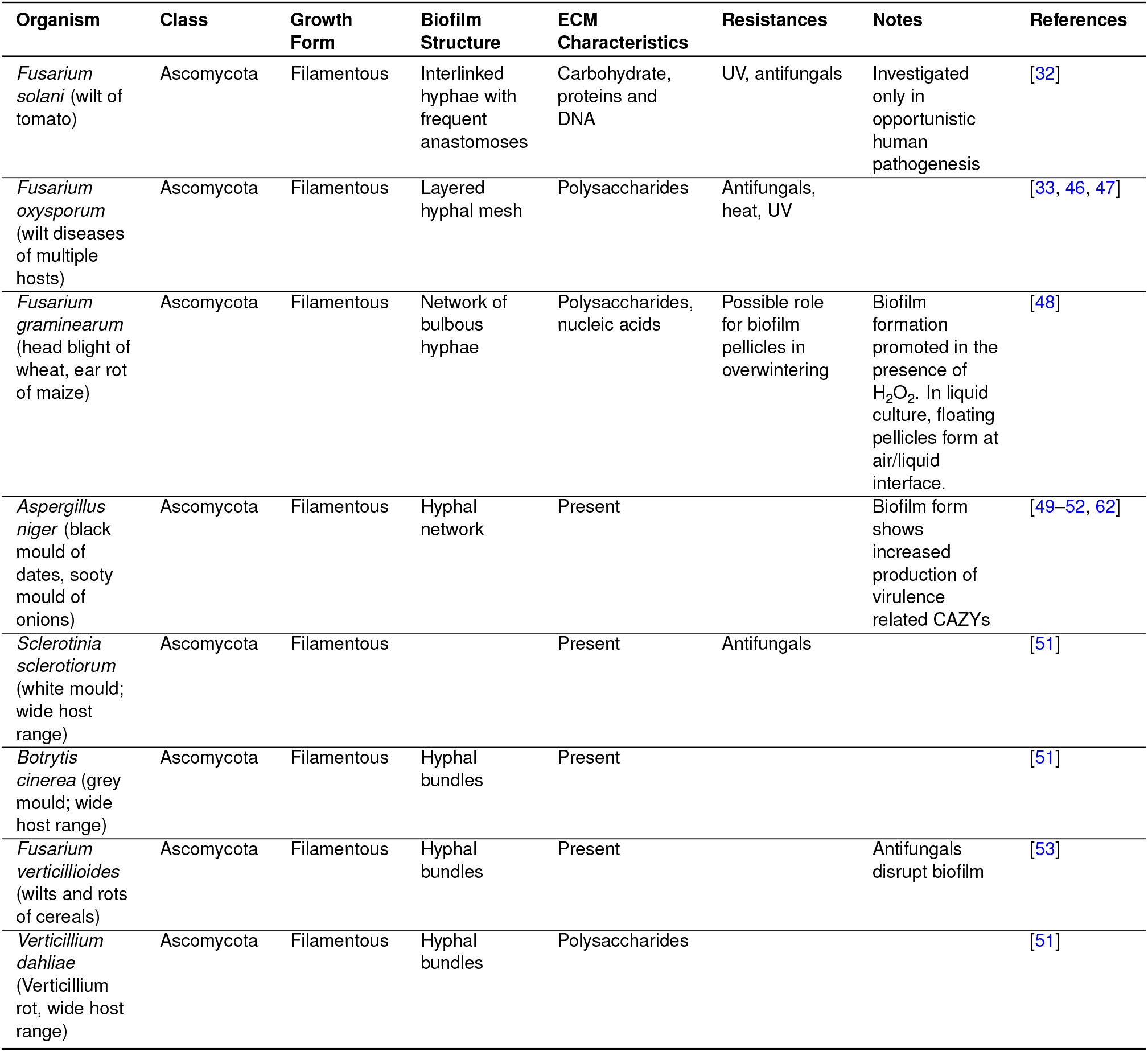
Known biofilm forming plant-pathogenic fungi show a number of shared features. Literature searches were performed using Web of Science, NCBI and Google Scholar databases with keywords ‘fungal’, ‘biofilm’ and ‘plant’ and information concerning the structure and properties of these biofilms summarised.

### Biofilm-like structures in *Z. tritici* are attached to their substrate

In order to determine whether biofilm-like growth in *Z. tritici* demonstrates the qualities associated with biofilms in other plant pathogenic fungi, we first investigated the attachment of cells to a hydrophobic surface using a washing and crystal violet staining assay. Biofilms in fungi have been shown to be dependent on carbon source[33, 63]; similarly, plant-derived nutrients such as sucrose[64] and the cutin monomer 16-hydroxyhexdecanoic acid[65] are known to alter fungal growth form. Therefore, a range of media and carbon sources were used in this assay. Stained cells adhering to the substrate were then visualised (Fig. 2).

**Figure 2.**
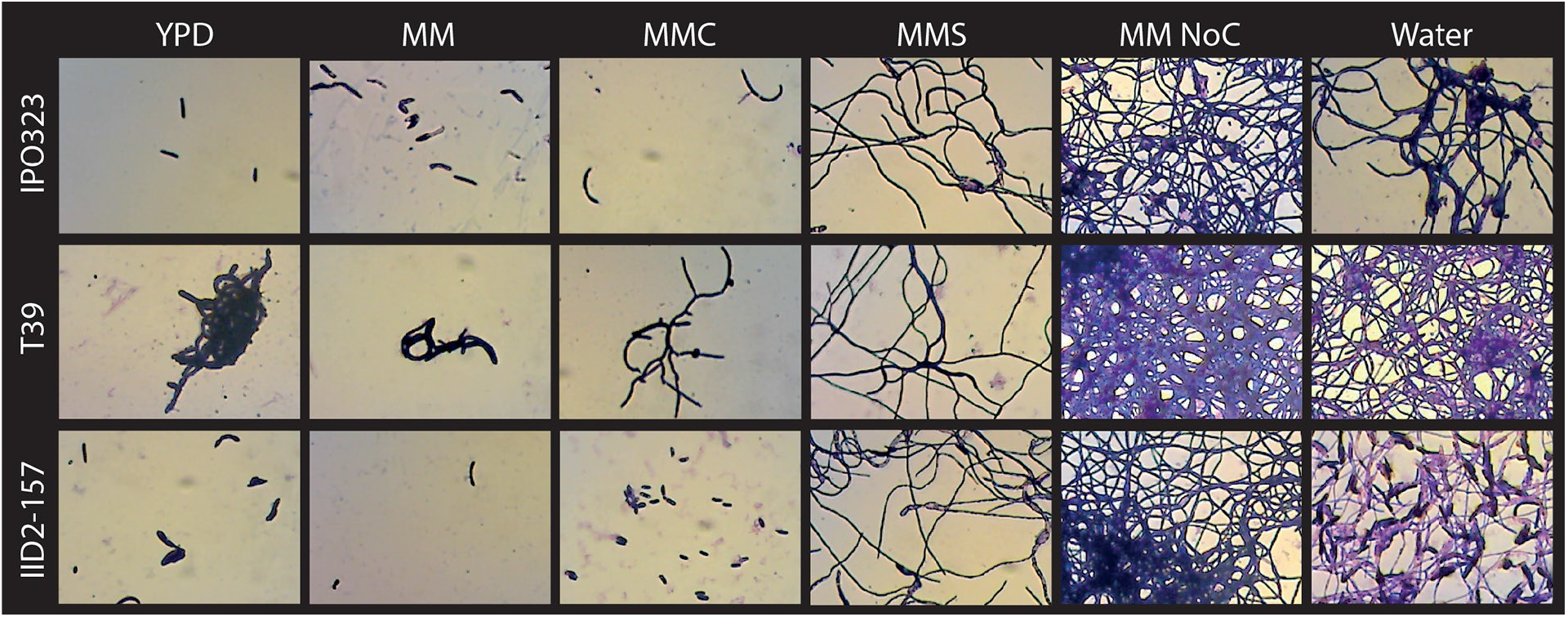
Adhesion of cells to substrate during *Z. tritici* growth in various media. *Z. tritici* isolates IPO323, T39 and IID2-157 were inoculated into YPD and MM, MMC, MMS, MM NoC or water in microtitre plates and incubated for 10 days at 20°C. Plates were then washed in sterile water to remove pelagic cells and adhered cells stained with crystal violet for observation.

There were clear differences between fungal growth form and degree of adhesion between the media. Neither complete medium, YPD (yeast peptone dextrose) or MM (minimal media) induced adhesion in any isolate, with few cells visible after washing and staining. The addition of the 16-hydroxyhexadecanoic acid to MM (MMC) did not appear to increase adhesion, although in isolate T39 it appeared to induce hyphal growth. The replacement of glucose in MM with sucrose (MMS) led to extensive hyphal growth and increased attachment to the substrate. Both growth in MM lacking any carbon source (MM NoC) and in water, however, led to very extensive attachment of hyphae and blastospores, forming a mat-like structure with bundles of blastospores connected by hyphae (Fig. 2), similar to structures observed *in planta* (Fig. 1). In addition, the fungal cells in these ‘mats’ appeared to be surrounded by and embedded in an extracellular matrix (ECM), another key feature of biofilms. This was particularly apparent in isolate T39 (Fig. 2), where channels in the ECM similar to those reported in other filamentous fungi[29, 30] were visible in the structures formed on MM NoC.

### Biofilm formation is medium and isolate-dependent

To quantify the observed differences in biofilm formation, adhesion and ECM production, we grew *Z. tritici* in media that produced visually different results in the previously assay: YPD, MMS and MM NoC. We then carried out a variant of the crystal violet assay for biofilms described by O’Toole[66]. This allowed the separate quantification of the attached stained cells and ECM (Fig. 3). When corrected for growth differences, both media type and isolate are significant predictors of both attachment of cells to the substrate and ECM production (2-way ANOVAs. Cells: medium, *p* = 0.0049; isolate, *p <* 0.0005. ECM: medium, *p* = 0.038; isolate, *p <* 0.0001). Tukey’s HSD tests indicate that differences between media were mainly driven by large increases in both cell attachment and ECM production in MM NoC, while isolate differences were due to similar increases in MMS for isolate IID2-157 only.

**Figure 3.**
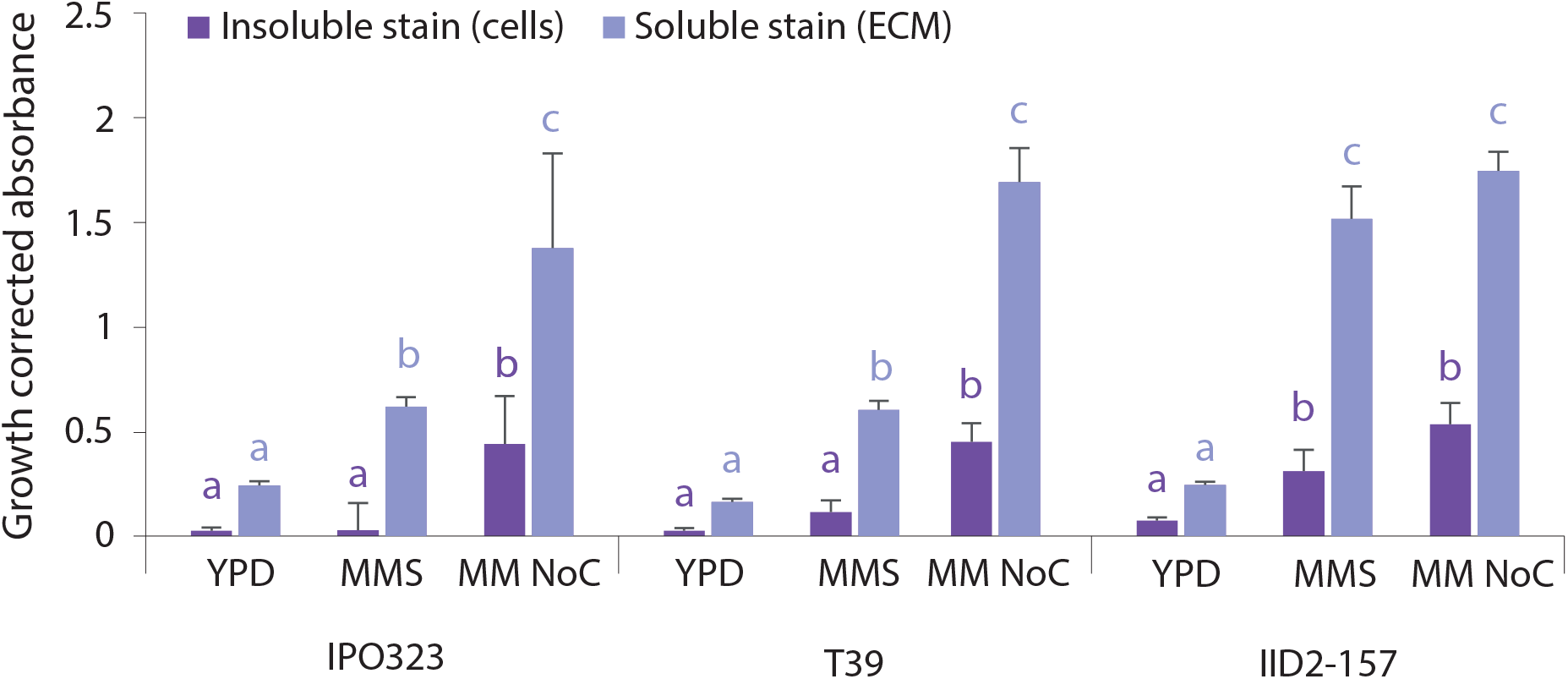
Low carbon conditions increase the key biofilm features of attachment and ECM production. *Z. tritici* isolates IPO323, T39 and IID2-157 were inoculated into YPD, MMS, MM NoC or water in microtitre plates and incubated for 10 days at 20°C. Plates were then washed in sterile water to remove pelagic cells and a modified crystal violet (CV) assay was used to stain and to differentiate between adhered cells (dark purple bars) and ECM (pale purple bars). Media-& isolate-dependent differences in growth-corrected OD550 values for adhered cells and ECM were assessed *via* two-way ANOVAs with Tukey’s simultaneous comparisons. Different letters above bars indicate significantly different in growth-corrected OD550 values (Cells: *p* = 0.0049; ECM: *p <* 0.0005) and isolates (Cells: *p <* 0.0005; ECM *p* = 0.005). Values are means of two independent experiments and error bars show SE.

### ECM is visible in scanning electron micrographs of cells grown in biofilm-promoting conditions

In order to confirm that the crystal violet stained non-cellular material associated with cells grown in MM NoC or water (Fig. 3) represented an ECM in which the *Z. tritici* cells were embedded in a biofilm, we used scanning electron microscopy (SEM) to visualise any material between cells or acting to attach cells to the substrate (Fig. 4). Polythene sheets were submerged in YPD or MM NoC inoculated with *Z. tritici* IPO323 at 10^5^ cfu mL^−1^ and incubated for ten days. After washing to remove pelagic cells, cells adhering to polythene pieces were chemically fixed and visualised. Cells grown in YPD were not abundant on the polythene pieces, but those present (Fig. 4A,B) had the expected cylindrical appearance, clearly visible septae, and showed no sign of any substance attaching them to the polythene substrate or to each other. However, cells grown in MM NoC showed a substance coating the spaces between the cells (Fig. 4C) and covering the cells themselves, obscuring the septae and extending projections which appeared to anchor the cell to the polythene substrate (Fig. 4D,E). In places, these projections extended and overlapped to form a web, binding the cell to the substrate (Fig. 4F). This web was, in places, very extensive, covering the substrate for more than a cell’s width beyond the cell boundary, with projections several times that in length. This web of extracellular material gave the cells a softer, less distinct edge when viewed at lower magnification (Fig. 4C), but was revealed at high magnification to be a matrix of criss-crossed filaments. Areas with higher cell density showed similar strands and webs linking the cells together (Fig. 4G,H). In areas of contiguous biofilm-like growth, these could be seen to coalesce into a matrix in which the cells were completely embedded (Fig. 4I). Pores or channels were visible within this matrix (Fig. 4H,I).

**Figure 4.**
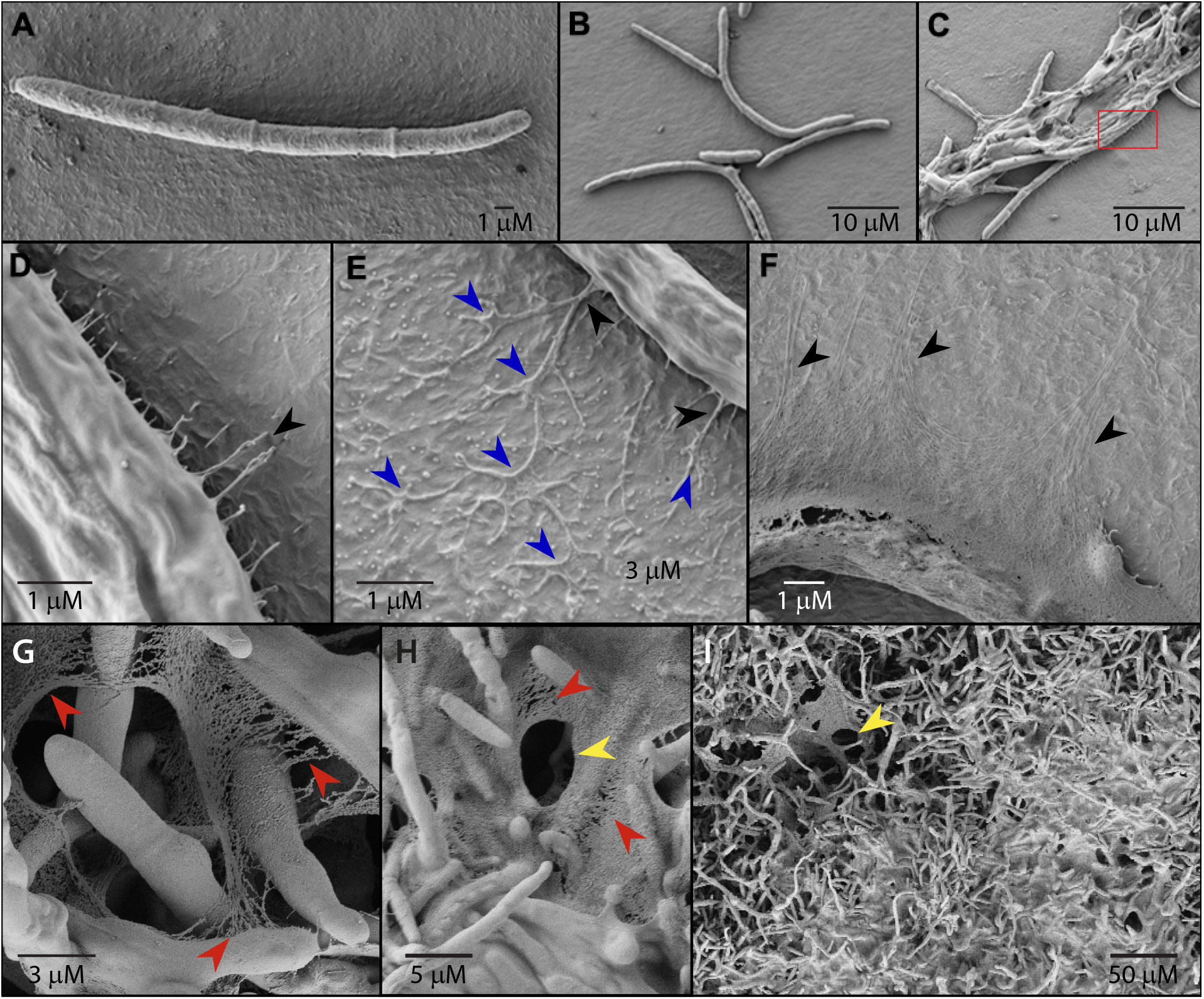
ECM production by *Z. tritici* cells. Cells were grown on pieces of polythene sheeting submerged in either YPD or MM NoC. 10 day old biofilms were viewed by scanning electron microscopy. In YPD, individual cells are seen, with clear differentiation between the cell edge and the substrate (A, B). By contrast, cells grown in MM NoC are seen aggregated and their boundaries with the substrate appear less distinct (C). High magnification images of these boundaries reveal projections (black arrows) from the fungal cells (D; magnified from red box in C) which develop into webbed structures adhering to the substrate (E; blue arrows). These webs can extend several times the width of the cell as a continuous sheet (F) which provides an explanation for appearance of the interface between these cells and the substrate at lower magnification. As biofilms develop and cells begin to aggregate, similar webs and strands join them (G, H; red arrows) and coalesce to form a continuous matrix in which the cells are embedded (H, I). Within the densest areas of this matrix, pores or channels are seen (H, I; yellow arrows).

### *Z. tritici* can produce multiple types of biofilm

Crystal violet staining and microscopy indicated that when grown in YPD, *Z. tritici* does not produce a biofilm that adheres to the substrate. However, when grown in 96-well microtitre plates, *Z. tritici* produced a visible, melanised ring of cells around the edge of the well at the air-water interface. This growth was not captured by OD readings due to its peripheral position in the wells, and was easily detached, meaning that it was not captured by the crystal violet assay. Indeed, since this assay measures attachment of cells and ECM to their substrate, this growth form should not be captured by it. However, biofilms can be formed at air-water interfaces [14, 48, 67] and in some definitions, attachment of cells to each other, rather than to a substrate, may be considered to create a biofilm[14, 68]. In the light of this, we decided to investigate this possible additional type of biofilm in *Z. tritici* IPO323. Blastospores were inoculated into YPD in petri dishes and allowed to grow for ten days, after which we observed a melanised film at the air-medium interface. In addition, we discovered a similar film of cells at the bottom of the media, which was not melanised and not attached to the petri dish surface. Both films were sampled and fixed for SEM visulisation (Fig. 5). Both films were composed of interwoven hyphae, with some blastospores also visible, and both showed extensive ECM (Fig. 5A, B). Films formed at the air-liquid interface showed chlamydospore production (Fig. 5A) and, when viewed at lower magnification, were revealed to have a distinctive structure of interconnected, roughly spherical masses of cells (Fig. 5C). Cells that remained pelagic in the culture medium showed no ECM and no interconnectivity (Fig. 5D).

**Figure 5.**
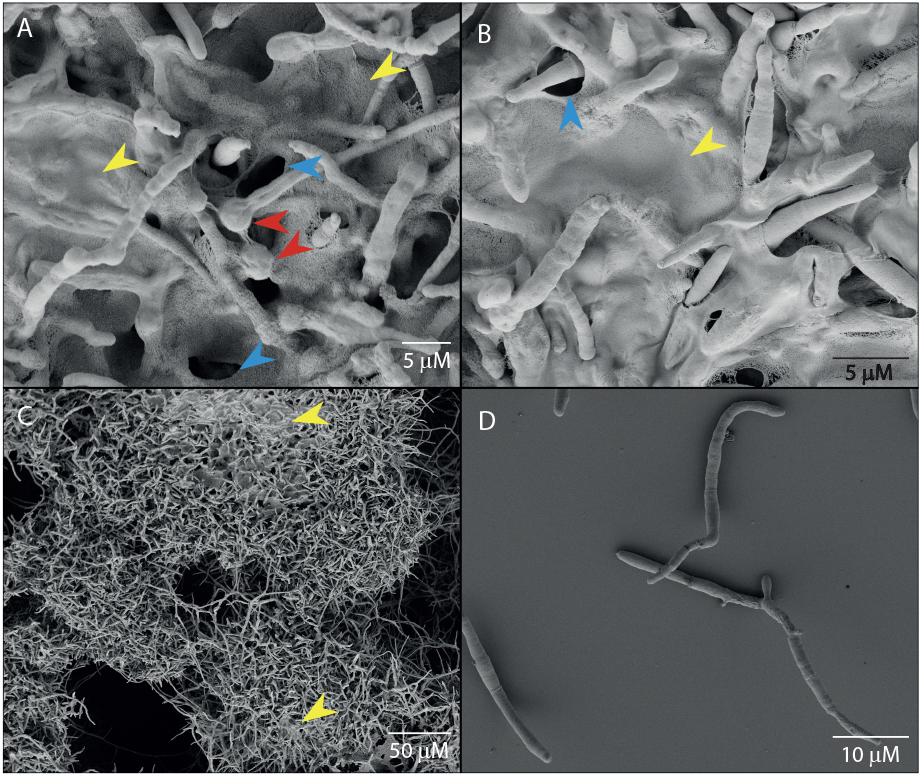
*Z. tritici* IPO323 produces two distinct forms of non-adherent biofilms when grown in static YPD. *Z. tritici* IPO323 blastospores were inoculated into YPD and maintained in static culture for 10 days. A melanised biofilm formed at the air-liquid interface (A, C) and a second non-adherent film formed at the bottom of the medium. Both films were composed of interconnected cells and could be harvested as a single mass. Scanning electron micrographs show: A. a portion of the air-liquid interface biofilm, showing hyphae and blastospores embedded in ECM (yellow arrows) with channels (blue arrows) and chlamydospores (red arrows) visible; B. a portion of the submerged non-adherent biofilm, showing hyphae and blastospores embedded in extensive ECM (yellow arrows) with channels (blue arrows); C. a lower magnification image of the air-liquid interface biofilm, showing the structure of interconnected masses of cells and the densest areas of ECM (yellow arrows); D, pelagic cells from the same media, showing no ECM or cell-cell adherence.

### Biofilm formation follows a set of predictable steps

SEM images of cells grown on polythene sheeting in MM NoC revealed that a predictable set of steps occurs during in biofilm formation. Single blastospores with no association to the polythene could be seen (Fig. 6A), which resemble pelagic cells. Once these are attached to the polythene, budding and elongation produces ‘tangled’ structures (Fig. 6B). Either these structures or individual blastospores can also act to ‘collect’ other blastospores which settle against them from the medium, forming bundles of aligned cells (Fig. 6C). As cells continue to grow, these ‘bundles’ and ‘tangles’ form denser structures (Fig. 6D) with visible ECM aiding adherence both between cells and to the substrate. At the periphery of these dense structures, hyphae grow outwards across the substrate. These hyphae then serve as foci for the further aggregation of cells (Fig. 6E). Aggregating cells then become interconnected via strands or webs of ECM, which can coalesce to completely encase neighbouring cells (Fig. 6F). The most extensive and mature biofilms (Fig. 6G) also show these strands or continuous areas of matrix encasing cells. In mature biofilms, the densest areas of ECM show evidence of blastospore production being the dominant growth form (Fig. 6G). This produces a ‘mat’ of blastospores that is similar to that seen *in planta* (Fig. 1).

**Figure 6.**
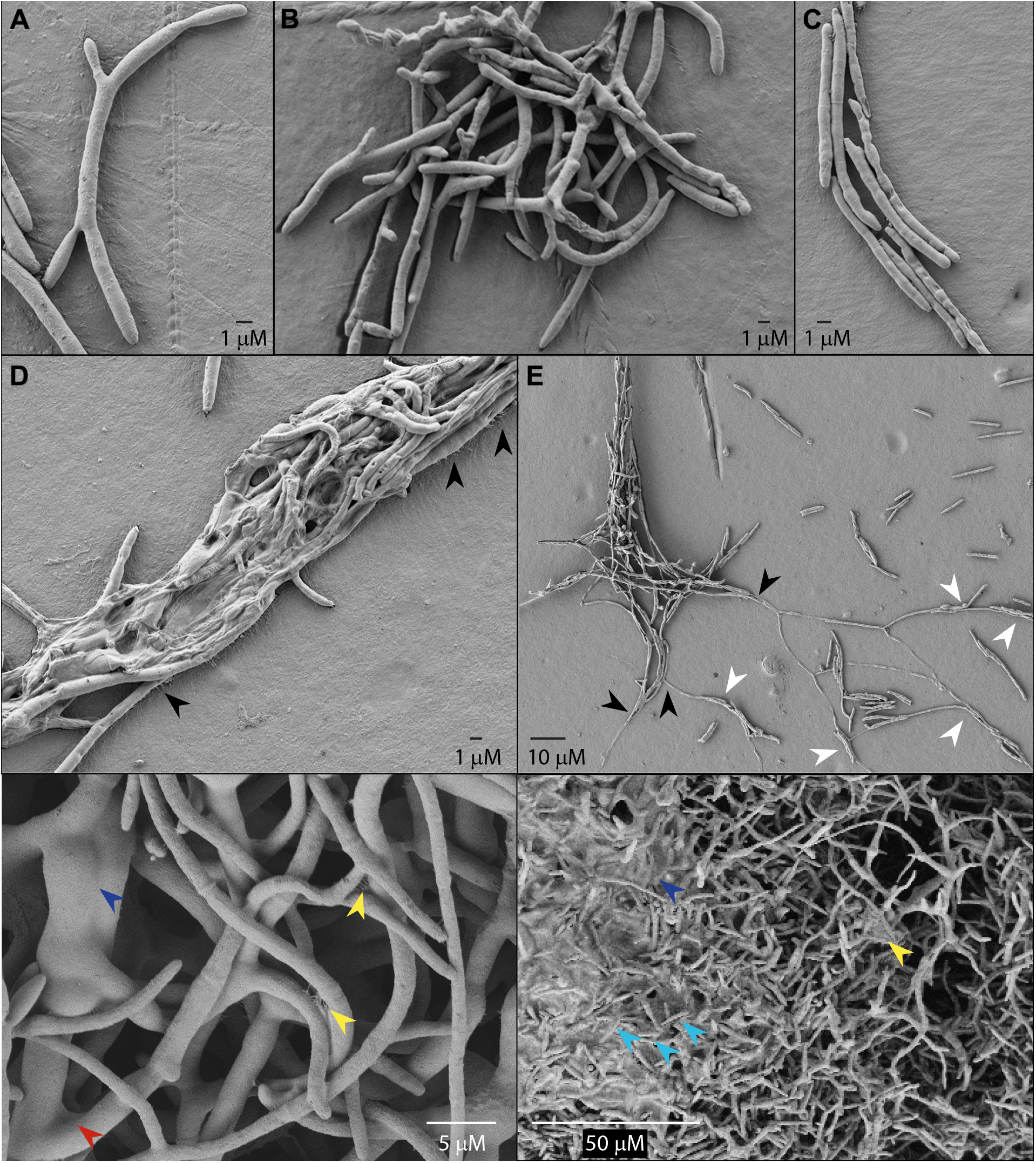
Development of biofilm structures on polythene by *Z. tritici* IPO323. Cells were grown for 10 days on pieces of polythene sheeting submerged in MM NoC. Resultant biofilms were viewed by scanning electron microscopy. Attached fungal cells/structures could be seen alone (A), grouped (B,C), or in various stages of biofilm development, allowing inferences to be drawn about the probable sequence of events involved in biofilm development. These images suggest that individual blastospores initially aggregate into tangled structures (B) or ‘bundles’ (C). Tangled structures appear to arise through growth, whereas bundles of aligned cells could be the simple result of cells settling against one another. These structures become more dense as cells continue to grow (D), and ECM production begins to be visible as strands connecting cells to their substrate (arrows, D). Cells at the edges of these denser structures germinate to form hyphae (black arrows, E), around which new aggregations of cells begin to form (white arrows, E). Aggregating cells then become interconnected *via* strands (yellow arrows, F) or webs (red arrow, F) of ECM, which can coalesce to completely encase neighbouring cells (blue arrow, F). The most extensive and mature biofilms (G) also show these strands (yellow arrows) or continuous areas of matrix encasing cells (blue arrow). In mature biofilms, the densest areas of ECM show evidence of blastosporulation (cyan arrows, G).

**Figure 7.**
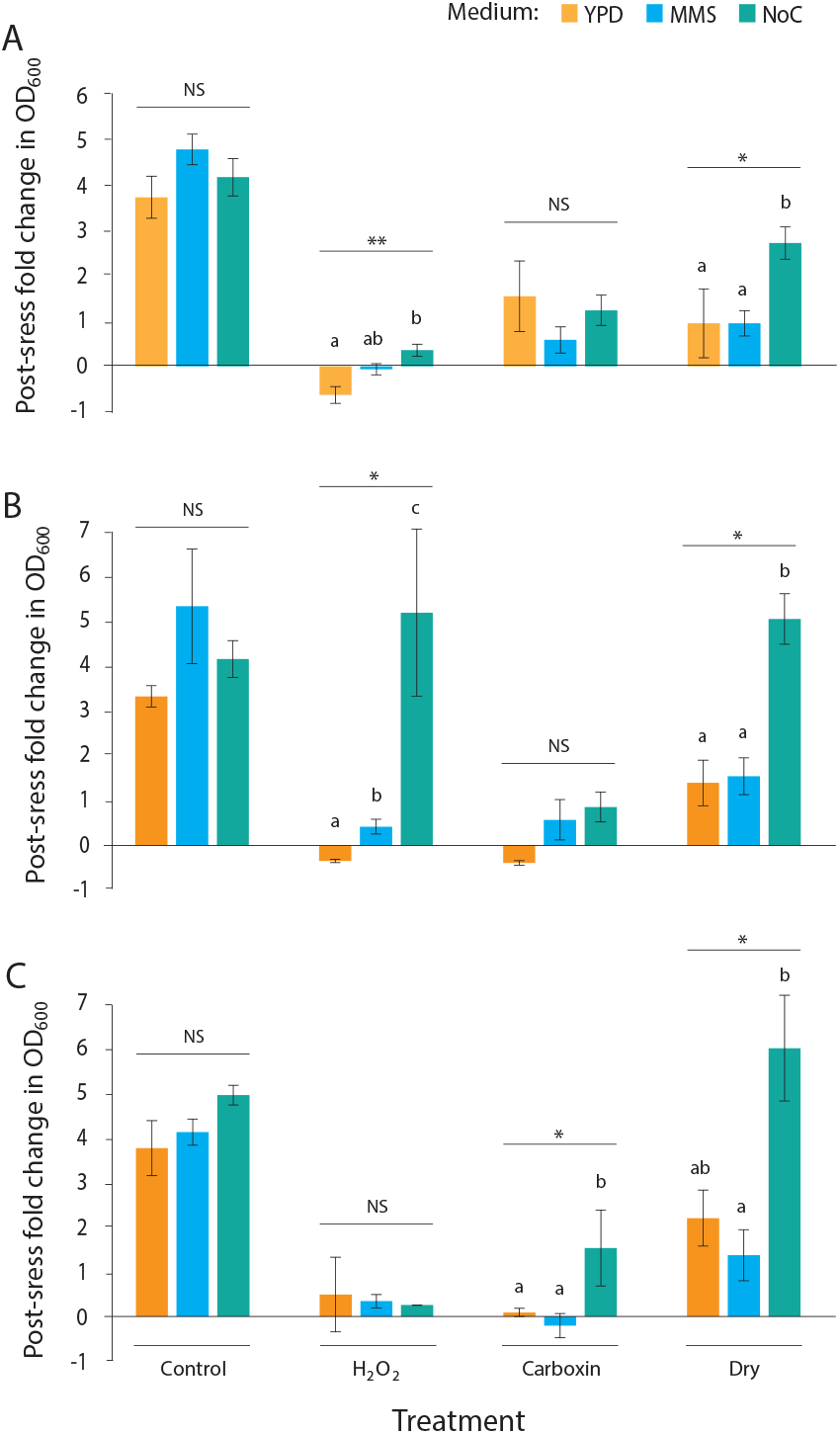
10 day old *Z. tritici* cultures show increased stress resistance when grown in a biofilm-inducing medium. *Z. tritici* cell suspensions (A: IPO323, B: T39 and C: IID2-157) were inoculated into either YPD, MM or MM NoC in microtiter plates. Plates were maintained for 10 days at 20°C. Cultures were then exposed to carboxin (47 µg mL^−1^), H_2_O_2_ or drying stress. Plates were then washed three times in sterile water to remove pelagic cells and stressors. 2 ml of fresh YPD was added to each well and OD_600_ values measured. Plates were then incubated for 5 days at 20°C. Increases in OD_600_ over this time period were calculated and are expressed as a % of the original growth in each medium during the 10 days prior to stress treatment, in order to control for differences in initial inoculum between media. Values are means of three independent experiments and error bars show SE. Asterisks represent significant differences in post-stress growth between media (ANOVAs - Controls: IPO323, *p* = 0.358; T39, *p* = 0.190; IID2-157, *p* = 0.404. H_2_O_2_: IPO323, *p* = 0.003; T39, *p* = 0.026; IID2-157, *p* = 0.170. Carboxin: IPO323, *p* = 0.378; T39, *p* = 0.135; IID2-157, *p* = 0.038. Drying: IPO323, *p* = 0.041; T39, *p* = 0.014; IID2-157, *p* = 0.029.). Different letters above bars represent significant differences in Tukey’s simultaneous comparisons.

### Biofilm formation leads to increased tolerance to elevated temperature and drying stresses, as well as resistance to the fungicide carboxin

In addition to adhesion and ECM production, a key feature of biofilms is their increased ability to withstand a variety of stressors. In order to determine whether *Z. tritici* biofilms show this property, 10-day old putative *Z. tritici* biofilms grown on polythene sheets were assessed for their ability to survive drying at 28°C, exposure to carboxin or exposure to hydrogen peroxide for four hours. Subsequent growth in fresh media added to washed biofilms was used as a measure of stress resistance, when compared to growth from unstressed washed biofilms. Initial biofilms were grown in either biofilm-promoting (MM NoC) media, MMS or YPD, and subsequent growth expressed as % change in OD to correct for media-induced differences in amount of biofilm available to seed the fresh media. All isolates showed an increase in growth following drying when pre-grown in the biofilm-promoting MM NoC (One-way ANOVAs - IPO323: *p*=0.041; T39: *p*=0.014; IID2-157: *p*=0.029), suggesting that the ability of cells to survive drying was higher in the biofilm. Two isolates showed media-dependent growth after treatment with H_2_O_2_. IPO323 showed little post-H_2_O_2_ growth in any medium, but post-H_2_O_2_ growth was significantly higher following pre-growth in MM NoC than in YPD (ANOVA; *p* = 0.003). T39 showed the same pattern (ANOVA; P = 0.026), but with extensive post-H_2_O_2_ growth following pre-growth in MM NoC. There were no media-dependent differences in post-H_2_O_2_ growth for IID2-157 (ANOVA; *p* = 0.170) Only IID2-157 showed media-dependent differences in growth following carboxin treatment, with greater growth in cells pre-grown in MM NoC (ANOVAs: IPO323, *p*=0.378; T39, *p*=0.135; IID2-157, *p*=0.038).

### Cells in biofilms show increased expression of some genes involved in adhesion and ECM production in other fungal biofilms

To determine whether genes known to be important for biofilm formation in other fungal species showed up- or down-regulation in *Z. tritici* biofilms, we carried out qRT-PCR of selected genes of interest (Fig. 8). Three investigated genes showed signficant upregulation in the YPD air-liquid interface biofilm: ZTRI_1.1499, ZTRI_-4.24 and ZTRI_9.291, with the greatest upregulation (>25x) seen in ZTRI_4.24. By contrast, ZTRI_3.465 was significantly downregulated in the YPD-air-liquid interface biofilm and ZTRI_1.1705 transcripts could not be detected at all in these cells. Expression patterns in the adhered, submerged MM NoC biofilms were somewhat different, with significant upregulation seen again in ZTRI_4.24, but not this time in ZTRI_1.1499 or ZTRI_9.291. ZTRI_1.1499 was in fact significantly downregulated in NoC biofilms while ZTRI_1.1705, undetected in YPD air-liquid interface biofilms, was significantly upregulated in NoC biofilms.

**Figure 8.**
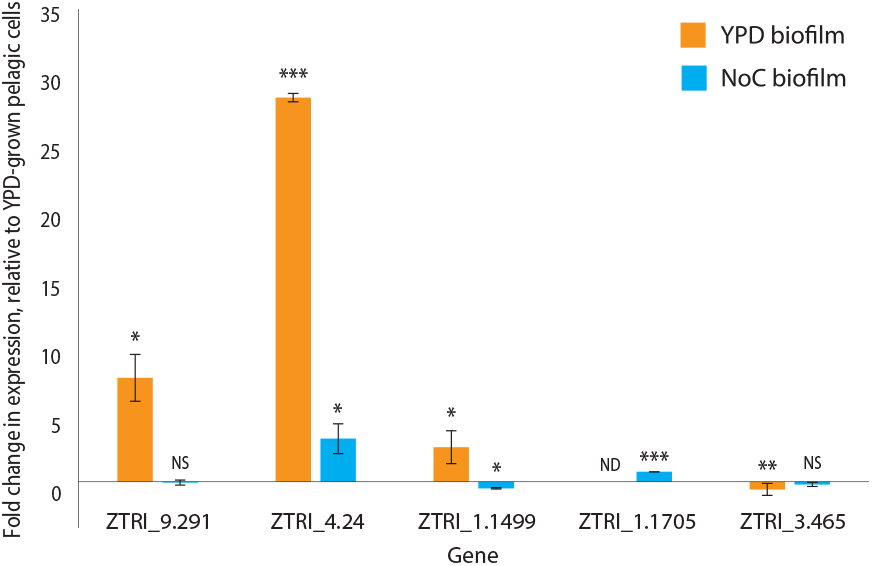
Expression of biofilm-related genes is distinct in both NoC- and YPD-grown biofilms, compared to pelagic cells. Pelagic IPO323 cells were harvested from shaken YPD cultures and biofilm IPO323 cells from static YDP or MM NoC cultures. RNA was extracted and qRT-PCR was performed to determine the expression of five biofilm-related genes of interest in biofilm cells from each media, relative to that in the YPD-grown pelagic cells. Fold change in expression is shown for the biofilm cells from both media, relative to expression in YPD-grown pelagic cells and normalised against the geometric mean expression of three separate reference genes (ELF1-*α*, UBC1 and *β*-tubulin) in each condition. Values are means and error bars show SE across three biological repeat samples of each growth condition, each containing three technical replicates. Significant change in gene expression relative to YPD-grown pelagic cells is represented by asterisks: * < 0.05; ** < 0.01; *** < 0.001. Significance determined using *t* -tests comparing sample means to 1 (= no change).

## Discussion

In this work, we have investigated the possibility that *Zymoseptoria tritici* is capable of biofilm formation. Initially, we observed biofilm-like growth *in planta* in two apparently very different interactions: isolate T39 on Galaxie, an almost entirely avirulent interaction where very few pycnidia are produced [59] and isolate IID2-157 on Volcani, a highly virulent interaction resulting in high-density pycnidiation. In both cases, we observed matlike structures formed mainly from short, blastosporelike cells, connected by hyphae. We interpret these cells as the result of budding on the leaf surface, as previously reported in *Z. tritici* by Francisco *et al*., 2019[69]. This biofilm-like phenotype is not seen in the commonly studied reference isolate IPO323 on susceptible wheat. For this reason, we hypothesised that our observations might represent a previously unobserved biofilmforming ability in certain isolates of *Z. tritici*. Review of the information available concerning biofilm formation in filamentous or dimorphic fungal plant pathogens showed that biofilm-formation on the host has a small number of precedents, mostly in *Fusarium sp*.. The precise structure of these biofilms varies, even among species in the same genus, with *Fusarium* species producing interlinked hyphal networks (*F. solani*[32]), layered hyphal mesh (*F. oxysporum*[33]), hyphal bundles (*F. verticillioides*[53]) or a network of hyphae with a bulbous morphology specific to ROS-stressed cells in biofilms (*F. graminearum*[48]). This variability reflects that seen in fungal biofilms more generally, where biofilms may consist of yeast-like cells, hyphae, or a mixture of these morphologies [20, 46, 48]. Given this variability, our *in planta* observations are consistent with a *Z. tritici* biofilm. Hyphal, yeast-like and mixed fungal biofilms share common properties, which are in turn shared with those of the better studied bacterial biofilms. These are: attachment of cells to a substrate; production of an extra-cellular matrix (ECM) and increased survival under stress [15, 17, 22]. For fungal plant pathogens, the stresses to which biofilm growth increased tolerance included heat, UV and ROS (Table 1). We adopted these commonalities as criteria which *Z. tritici* biofilm-like growth should meet to be considered a true biofilm: 1) attachment to a substrate; 2) ECM production; 3) increased stress tolerance.

To test whether our putative biofilms met these criteria we assessed biofilms produced *in vitro*. We hypothesised that *Z. tritici* would be most likely to produce similar growth to that observed on the leaf surface when grown in conditions that mimicked that environment. We therefore grew isolates T39 and IID2-157, plus reference isolate IPO323, in a range of media. We visualised putative biofilms using crystal violet staining. Staining involves multiple washing steps, such that the visualised fungal material is that which is adhered to the substrate. With YPD as a high-nutrient control, we compared minimal medium (MM) to sucrose minimal medium (MMS), MM + 16-hydroxyhexadecanoic acid (MMC), carbon-free minimal medium (MM NoC) and water. Sucrose is commonly detected on the surface of cereal leaves, and 16-hydroxyhexadecanoic acid is a cutin monomer found in their waxy cuticle layer[65]. Thus, these two media potentially resemble the wheat leaf surface more than MM. Both MM NoC and water were included in order to reflect the low-nutrient availability environment of the leaf surface. There were clear visual differences between the putative biofilm morphologies in the different media. In YPD, MM and MMC we observed few stained cells attached to their substrate. These cells appeared to be predominantly in the yeast-like/blastospore growth form. MMS promoted the hyphal growth form, with increased, but still low, attachment. This is in line with the previous report that sucrose promotes the switch to hyphal growth[64]. By contrast, growth of *Z. tritici* in both MM NoC and water led to extensive attachment to the substrate by a mixture of hyphal and yeast-like growth, apparently surrounded by an extracellular matrix. These observations were consistent across the three *Z. tritici* isolates. These results indicate that both blastospores and hyphal growth forms are capable of attachment to the substrate. The densest biofilms contained both cell types, and ECM appeared as a substance filling the gaps between interwoven hyphae. The three media that showed increased attachment of cells (MMS, MM NoC and Water) were all media that promoted hyphal growth, suggesting the possibility that hyphae are important for the formation of a stable biofilm formation, as seen in other dimorphic fungi such as *Candida*[70, 71]. The apparently random or unstructured distribution of blastospores at locations with the network of hyphae, however, indicates that the layered structure of hyphae built on a foundation of attached yeastlike cells that is characteristic of *Candida* biofilms[17] is not seen in *Z. tritici*. The most extensive biofilms were formed in the no carbon or no nutrient media, MM NoC and water. This indicates that *Z. tritici* biofilm formation is induced or increased under carbon limitation. This is in contrast to *Fusarium* and *Cryptococcus* biofilms, whose formation has been shown to be carbon dependent[33, 71]. However, *Z. tritici* blastospores have very extensive lipid stores, hypothesised to be important in sustaining growth during long periods on the leaf surface and *ex-planta*[60]. These stores may provide a substitue for external carbon during biofilm formation. It is further possible that *Z. tritici* biofilms contain hypoxic environments in which some cells have very low metabolic activity, as seen in other biofilms[17, 34]. In this case, biofilm formation might be an adaptation to low nutrient environments, allowing the fungus to reduce its metabolic rate and survive in the absence of sufficient carbon sources. This might be particularly useful on resistant or non-host plants, where leaf penetration and access to apoplastic nutrients is not possible. We hypothesise that the groups of blastospores seen in these biofilms might therefore have a role in dispersal to new environments *via* rain-splash, since dispersal mechanisms are a commonly observed feature of mature biofilms[14, 15, 48].

To quantify these observed differences in cell adherence and ECM production, we used a crystal violet (CV) assay conducted in 6-well microtiter plates to compare biofilm-inducing MM NoC to the hyphal-growth promoting MMS and to YPD controls. Ethanol-soluble and insoluble CV stain (staining the ECM and cells, respectively) could be differentiated, giving an estimate of the proportion of CV in ECM vs attached cells. This assay confirmed a significant increase in cell attachment and ECM production for all isolates under putative biofilm-inducing MM NoC conditions. MMS produced similar increases in attachment and ECM production for the isolate IID2-157, but in ECM production only for other isolates. Thus, we conclude that MM NoC is a biofilm- and ECM-inducing medium for *Z. tritici*. Scanning electron microscopy confirmed the presence of ECM surrounding cells grown on a polythene surface in MM NoC. ECM has a fibrous appearance and seems to attach fungal cells to the polythene, beginning as strands linking the cells and substrate before developing into web-like protrusions which form a mesh of overlapping strands and eventually a matrix in which mutliple cells are embedded. At present, we cannot comment on the composition of this ECM. Polysaccharides, proteins, and nucleic acids have all been detected in the ECM of fungal biofilms[29, 32, 48]; however, CV stains carbohydrates, lipids and DNA, so is not useful for differentiating between these. The fact the CV can be solubilised from *Z. tritici* ECM with ethanol could indicate that the ECM is porous and allows the solvent to access the dye. However, it is also possible that the ECM itself is partially or completely ethanol soluble, as would be the case if it were lipid-based. Further staining and analysis will be required to determine what the ECM consists of and how it is organised into the structures observed. We also observed a pattern of steps in the formation of *Z. tritici* biofilms, involving cell attachment to the substrate, followed by the formation of bundles of blastospores and ECM production. Cells would then germinate and hyphae ramify across the substrate, acting as foci for the further aggregation of blastospores. This process is similar to that seen during biofilm development in other dimorphic fungi, such as *Candida*, the first step of biofilm formation is the attachment of yeast-like cells to a substrate, followed by the formation of a yeast-like cell layer from which the overlying hyphal layer originates[20, 27]. Mature biofilms showed cells fully embedded in ECM which not only linked them to the substrate, but to each other. This ECM showed channels or pores throughout. Mature biofilms showed many yeast-like cells on their surfaces, suggesting that blastosporulation was occurring. The biofilms that we observed at the air-liquid interface of static YPD cultures were distinctive in that they were melanised, and both these and the submerged biofilms seen in static YPD cultures were not adherent to any substrate, although the cells within them were adhered to one another by ECM, allowing the biofilm to retain integrity when pipetted or lifted on a glass coverslip. Visualisation by SEM showed that air-liquid interface biofilms were composed of interconnected cell masses embedded in ECM. Clamydospores were present in these masses, but were not seen in other biofilms.

The final criterion to consider the structures observed in *Z. tritici* to be ‘true’ biofilms was stress-resistance. To evaluate this, we exposed putative biofilms grown in MM NoC and control cultures grown in YPS or MMS to three different stressors: hydrogen peroxide, the fungicide carboxin, and a combination of 28°C and drying. These represent common stressors encountered during infection in the field: ROS-based plant defences, fungicides and heat/drying stress may all be encountered by *Z. tritici* prior to leaf penetration. For all isolates, growth in the biofilm-inducing MM NoC led to increased post-stress growth following heat and drying stress compared to YPD and MMS. Growth after H_2_O_2_ exposure was significantly greater in MM NoC than the other media for IPO323 and T39, whereas biofilm-inducing MM NoC only promoted growth following exposure to carboxin for the third isolates, IID2-157. From these results, it is clear that biofilm-inducing conditions lead to a more stress resistant growth form, as expected under the hypothesis that *Z. tritici* forms biofilms *in vitro* and on the wheat leaf. However, the degree of protection against each stress tested appears to be isolate dependent. This might reflect underlying differences in the isolates’ resistance to these stresses, irrespective of biofilm-formation, or differences in the structure of the biofilm, its penetrability for the different chemical stressors, or the composition or quantity of the ECM produced by different isolates.

Expression patterns of genes of interest were different between biofilm and pelagic cells, and there were also differences in their expression between the MM NoC, submerged, substrate-adhered biofilms and the biofilms found at the YPD air-liquid interface. ZTRI_4.24, a gene with orthologues in the ALS (agglutinin-like) gene family in *Candida albicans* (FungiDB[72]) and therefore likely to be involved in either cell-surface or cell-cell adhesion, was upregulated in both biofilms. However, the similarly annotated ZTRI_3.465 was downregulated in both biofilms. Interestingly, orthologues of both of these genes in *C. albicans* are annotated in FungiDB as induced in *C. albicans* spider biofilms but repressed in *C. albicans* rat-catheter biofilms[72]. This implies that different agglutinins may be used to mediate adhesion in different environments. Since the YPD biofilms were not adherent to the substrate, it is reasonable to hypothesise that ZTRI_4.24 may encode a protein used particularly in cell-cell adhesion. ZTRI_1.1705 another putative ALS-family protein - and ZTRI_1.1499 showed differential expression between the two types of biofilms. ZTRI_1.1499 is an orthologue of *C. albicans* Csh1. The CSH1 protein is implicated in cell surface hydrophobicity [73–75]. A similar function in *Z. tritici* would consistent with the upregulation we saw in the air-liquid interface YPD biofilm vs downregulation in the submerged NoC biofilm.

The greatest change in expression was seen for ZTRI_- 9.291, which showed >25-fold upregulation in the YPD biofilm and 4-fold upregulation in the NoC biofilms. This gene is implicated in the regulation of biofilm formation, but has been implicated in both negative[15, 76] and positive[77, 78] regulation of biofilm formation in *C. albicans*, as well as positive regulation of biofilm formation in the filamentous fungus *Aspergillus fumigatus*. Thus, our results are compatible with previous findings of a positive association between expression of orthologues of this gene and biofilm formation, although the mechanisms are unclear. It is notable that ZTRI_9.291 is upregulated in YPD-grown, but not MM NoC-grown biofilms. Since alcohol dehydrogenases are part of the primary carbon metabolism, and therefore likely to be differentially regulated under low carbon growth conditions, this may indicate differential strategies for forming biofilms and their extra-cellular matrices depending on nutrient availability. Interestingly, Adh1 has also been defined as a core stress-response gene in *C. albicans*[77].

Taken together, these results indicate that *Z. tritici* is able to produce structures which meet the definition of a biofilm. On a substrate such as polythene sheeting or the base of a microtiter plate well, *Z. tritici* adopts a specific biofilm growth form. This consists of dense areas of blastospores, connected by hyphae and attached to their substrate by an extra-cellular matrix. The hyphae are often arranged in bundles within a dense interwoven network of layered hyphae embedded in ECM which appears to include water channels similar to those reported in other fungal biofilms present a structure similar to that seen in other filamentous plant pathogen biofilms[29, 36]. In particular, the images we obtained are strikingly similar to those reported for *Fusarium solani* by Córdova-Alcántara *et al*. (2019)[32]. We saw similarly structured biofilms at the YPD air-liquid interface, with the addition of visible melanisation and increased expression of a gene that may be involved in cell surface hydrophibicity.

These biofilms show reduced sensitivity to combined temperature and drying stresses, as well as increased fungicide and ROS resistance in some isolates. Based on the observed phenotype of the isolates T39 and IID2-157 during epiphytic growth on resistant and susceptible wheat, respectively, we propose that biofilm formation occurs *in planta* during the epiphytic phase of the *Z. tritici*-wheat interaction[54]. Biofilm formation *in planta* is not specific to either resistant or susceptible wheat (or, therefore, to avirulent or virulent fungal isolates), but is not seen in all wheat cultivar-fungal isolate combinations. To our knowledge, this biofilm phenotype has not been reported for the reference isolate, IPO323.

If biofilms are formed on the surface of wheat leaves under field conditions, then it is likely that they contribute to the ability of *Z. tritici* to survive stresses encountered during the infection of crops. Biofilms able to withstand drying stresses, UV and fungicides could promote fungal survival on the leaf surface either prior to leaf penetration or when on a resistant wheat or non-host plant. Since nutrient availability on the leaf surface is low, it is likely that reduced the metabolism of cells in biofilms could be particularly beneficial when *Z. tritici* is growing epiphytically on a non-host or resistant wheat cultivar, unable to access apoplastic nutrients. *Z. tritici* blastosporulation has been previously reported on the leaf surface[69]. Given the role of reversion to yeast-like growth for dispersal in some fungal biofilms[15, 48], it is reasonable to speculate that blastospores produced in biofilms might be dispersed to nearby host plants, for example *via* rain-splash.

The possibility of *Z. tritici* biofilms being adaptations for survival and dispersal means that they may have a role to play in maintaining fungal isolates in the environment – and thus in the gene pool of the fungus – in the absence of a host. While the longevity of *Z. tritici* biofilms in field conditions is not known, the ability to form biofilms may allow *Z. tritici* to survive between growing seasons and/or disperse *via* sub-optimal routes such as the leaves of non-hosts. This possibility has implications for our understanding of the evolutionary dynamics of stress tolerance in *Z. tritici*, including fungicide resistance, tolerance of host defences, survival under the stresses imposed by climate change, and more.

## Methods

### *Zymoseptoria tritici* isolates

The *Z. tritici* isolates used in this study were kindly provided by Prof Gert Kema and Prof Gero Steinberg. Isolates IPO323 and IID2-157 were isolated in the Netherlands and have been widely used in studies of *Z. tritici*, with IPO323 commonly regarded as the reference isolates for work on this fungus[80–82]. Isolate T39 was isolated in the USA, from North Dakota[80]. IPO323 and T39 were isolated from bread wheat; IID2-157 from durum wheat. Strains of each isolate expressing cytosolic GFP were provided by Dr Sreedhar Kilaru. In each case, the GFP construct was integrated into the *sdi1* locus according to the methodology published in Kilaru *et al*. (2015)[83].

### Wheat plants and inoculation of wheat with *Z. tritici*

The wheat cultivars used in this work were Galaxie (bread wheat) and Volcani (durum wheat). Galaxie is susceptible to isolate IPO323 and largely resistant to isolate T39; Volcani is susceptible to IID2-157. Wheat was grown on John Innes No. 2 compost in a growth chamber at 20°C, 80% RH, 12 h light. Inoculations were carried out on 14 day-old wheat seedlings. Fungal spore suspensions were filtered through two layers of sterile Miracloth and adjusted to 10^6^ cfu mL^−1^/ml in 0.1% (v/v) Tween-20. Fully expanded wheat leaves, marked at the base, were coated with spores suspensions using a paintbrush. Inoculated plants were returned to the growth chamber and covered with cloches for the first 72 h. Cloches were then removed, and plants maintained under the same conditions.

### Fungal growth conditions and media

*Z. tritici* isolates were cultured from 50% glycerol stocks maintained at -80°C by plating onto yeastpeptone-dextrose (YPD) agar. Five day old blastospores were scraped from these plates and resuspended in either YPD, sterile distilled water (SDW), minimal medium (MM: 6 g L^-1^ NaNO_3_; 0.52 g L^−1^ KCl; 0.52 g L^−1^ MgSo_4_; 1.52 g L^−1^ KH_2_PO_4_; mg L^−1^ ZnSO_4_; 1.1 mg L^−1^ H_3_BO_3_; 0.5 mg L^−1^ MnCl_2_; 0.5 mg L^−1^ FeSO_4_; 0.16 mg L^−1^ CoCl_2_; 0.16 mg L^−1^ CuSO_4_; 0.11 mg L^−1^ (NH_4_)_6_Mo_7_O_24_; 5 mg L^−1^ Na_4_EDTA; 10 g L^−1^ glucose[64]), or a variant of MM at 10^7^ cfu mL^−1^. MM variants were: sucrose minimal medium, MMS, in which glucose was replaced with 10 g L^−1^ sucrose; No carbon minimal medium, MM NoC, in which glucose was omitted and not replaced; cutin minimal medium, MMC, in which the cutin monomer 16-hydroxyhexadecanoic acid was add to complete MM to give a final concentration of 200 µM.

### Crystal Violet assay for ECM and adhered cells

To quantify the key biofilm attributes of ECM production and adherence of cells to substrates, we used a modified version of the crystal violet assay described by O’Toole[66]. *Z. tritici* blastospores were resuspended from agar plates and suspensions adjusted to 10^7^ cfu mL^−1^ using a haemocytometer. 200 µL of this suspension was added to 2 mL of growth media in 24-well microtiter plates. Plates were incubated for 10 days at 20°C to allow biofilm formation. OD_600_ (fungal growth) was then read using a Clariostar plate reader. Media and pelagic fungal growth were then decanted and plates washed 3x in sterile water. 2 mL of 0.1% (w/v) crystal violet (CV) solution was then added to each well and incubated at room temperature for 30 mins. CV was decanted and the plate washed 3x with water and allowed to dry for 2 h at 28°C before the addition of 2 mL ethanol to solubilise the CV. Plates were gently shaken for 15 min and OD550 (CV absorbance total) was then measured. Finally, the ethanol was transferred to a clean plate and another OD550 measurement taken (CV absorbance ECM). CV stained *Z. tritici* cells attached to the plate retain their purple colouration after 15 mins in ethanol, and remain attached to the plate; meanwhile, CV bound to ECM is solubilised by the ethanol. Thus, the second OD550 measurement, taken in a clean plate, allows the absorbance of 550 nm light by CV solubilised from stained ECM to be distinguished from that due to stained, attached cells.

### Light microscopy of crystal violet stained cells

To visualise fungal biofilms, 500 µl of a 10^7^ cfu mL^−1^ blastospore suspension was added to 5 mL of growth media in 6-well microtiter plates. Plates were incubated for 10 days at 20°C to allow biofilm formation. Media and pelagic fungal growth were then decanted and plates washed 3x in sterile water. 2 mL of 0.1% (w/v) crystal violet (CV) solution was then added to each well and incubated at room temperature for 30 mins. CV was decanted and the plate washed 3x with water and allowed to dry for 2 h at 28°C before viewing at 10 x magnification using a light microscope. Representative images were obtained using an details eyepiece camera.

### Confocal laser scanning microscopy of infected wheat leaves

To observe *Z. tritici* isolates growing on wheat leaves, samples of leaves inoculated with GFP-tagged strains of fungal isolates were mounted in 0.1% (v/v) phosphate buffered saline (PBS, pH 7) and if necrotic, stained with 5 µL 0.05% (w/v) propidium iodide (PI). Confocal microscopy was carried out using argon laser emission at 500 nm with detection in 600-630 nm (chlorophyll/PI, red) and 510–530 nm (GFP, green), using a Leica SP8 confocal microscope [84].

### Scanning Electron Microscopy (SEM)

For ultrastructural analysis, cells were inoculated onto small pieces of polythene sheeting submerged in either YPD or MM NoC and incubated for 10 days at 20°C to allow biofilms to develop on the polythene. After removal from the growth media, polythene pieces were washed three times in water to remove pelagicpelagic cells. Adhered cells were then fixed in 2% paraformaldehyde and 2% glutaraldehyde in 0.1M sodium cacodylate, pH 6.8 for 2 h at room temperature. To that effect, the polythene pieces were fixed onto the bottom of plastic petri dishes using adhesive tape applied to the edges of the plastic sheets, then carefully covered with the fixative. After 3 x 5 min washes in cacodylate buffer, the samples were post-fixed in 1% aqueous osmium tetroxide for 1 h at RT. Following 3 x 5 min washes in deionised water, cells were dehydrated in a graded ethanol series (30, 50, 70, 80, 90, 95% - 5 min per step, followed by 2 x 10 min in 100% ethanol) and subsequently incubated for 3 min in HMDS (Hexamethyldisilazane, Merck, Gillingham, UK) before quickly air drying the samples. Dried samples were mounted on aluminium stubs with the help of carbon tabs and coated with 10 nm gold-palladium (80/20) in a sputter coater (Q150T, Quorum, Lewes, UK) before imaging in a scanning electron microscope (Zeiss Gemini 500 SEM) operated at 1.5 kV and a SE2 detector.

### Biofilm stress tolerance assays

To determine whether biofilms showed altered survival under stress conditions, 200 µL of 10^7^ cfu mL^−1^ suspensions of all three isolates were inoculated into 2 mL of YPD or MM NoC in 24-well plates (2 wells per isolate-medium combination). Plates were incubated at 20°C for 10 days to allow biofilm formation. Plates were then subjected to one of four stress conditions (3 plates per stress condition): 1) Control: plates were maintained at 20°C for 4 h; 2) Carboxin: 2 µL of carboxin solution (47 mg mL^−1^ in DMSO) was added to each well and plates incubated for 4 h at 20°C; 3) H_2_O_2_: 250 µL of H_2_O_2_ (30% v/v in water) was added to each well and plates incubated for 4 h at 20°C; 4) Drying/heat: media was decanted and wells rinsed three times with SDW. Lids were then replaced with a single layer of sterile miracloth and plates incubated at for 4 h at 28°C, which was sufficient to evaporate all remaining SDW. For stresses 1-3, after the 4 h incubation, media was decanted and wells rinsed three times with SDW to remove any non-adhered cells, decanting each time. 2 mL of YPD was then added to each well for all plates and a further OD_600_ reading taken [OD_600_-stress]. Plates were then incubated for 4 days and a final OD_600_ reading taken [OD_600_-final]. Stress tolerance was then calculated per well as [OD_600_-final]-[OD_600_-stress]/[OD_600_-stress]. This calculation represents the net growth in fresh medium derived from the stressed biofilms as inoculum. Dividing by the [OD_600_-stress] reading for each well corrects for differences in the amount of biofilm available to act as inoculum, which is isolate and medium dependent. Thus, the final value calculated represents net growth per unit of inoculum and provides a proxy for the percentage of the original inoculum which remained alive following the application of the stress conditions.

### Investigation of YPD biofilms and pelagic cells

To investigate the biofilms formed in the rich culture medium YPD, 200 µL of 10^6^ cfu mL^−1^ blastospores were inoculated into 20 mL YPD broth in a petri dish and maintained at 20°C without shaking or disturbance for 10 days. Air-media interface and non-attached, submerged films were harvested by gentle pipetting, using a 1 mL pipette tip with the narrowest 15 µL cut off. Pelagic YPD-grown cells were harvested from 20 mL YPD cultures inoculated in the same way as biofilm cultures but incubated in 50 mL falcon tubes with 240 rpm shaking at 20°C for 10 days. Biofilms and pelagic cells were then transferred to ependorf tubes for staining with CV or fixing and preparation for SEM visualisation.

### RNA extraction from biofilms and qRT-PCR

To investigate gene expression, RNA was extracted from *Z. tritici* cells grown under biofilm-forming and control conditions. 500 µl of a 10^7^ cfu mL^−1^ suspension of IPO323 was inoculated into 30 mL of YPD or MM NoC in either petri dishes (biofilms) or 50 mL falcon tubes (pelagic cells). Plates were incubated at 20°C for 10 days to allow biofilm formation, while falcon tubes were incubated at the same temperature with 200 rpm shaking for ten days. Pelagic cells were harvested by centrifugation, supernatant was discarded and the cell pellet flash frozen in liquid nitrogen. Media and pelagic cells were decanted from petri dishes and biofilms rinsed once in fresh media to remove non-adherent cells. A liquid shear was then performed with 5 ml 0.05% (w/v) Tween-20. Detached cells were decanted into 15 ml falcon tubes and harvested by centrifugation. The supernatant was discarded, the cells pellet rinsed once in growth media, and flash frozen in liquid nitrogen. Cell pellets were then stored at -80°C until required.

To extract RNA from harvested cells, pellets were thawed on ice and resuspended in 1 ml TRIzol solution (Invitrogen, UK). Samples were transferred to 2 ml Safe-Lock microcentrifuge tubes and homogenised in a QIA-GEN TissueLyser II for 2 min at 28 Hz. From this point RNA was extracted following the TRIzol RNA extraction protocol as per manufacturer’s instructions. RNA concentration was measured and purity assessed using a Nanodrop-1000; integrity was checked by gel electrophoresis. RNA was normalised to the lowest sample concentration and cDNA libraries prepared using the QuantiTect® Reverse Transcription Kit (QIAGEN, UK) as per manufacturer’s instructions. cDNA integrity was also checked by gel electrophoresis.

Primers were designed using the Clone Manager software (Sci Ed). Genes of interest from the *Z. trtici* genome were chosen by homology to genes from *Candida albicans* or *Aspergillus fumigatus* known to be involved in biofilm formation. These were: ZTRI_1.1499, an orthologue of the *Candida albicans* gene *CSH1*[72], implicated in cell surface hydrophobicity[74, 75] and ECM production[73] and annotated in FungiDB[72] as being positively and negatively regulated during biofilm formation in *C. albicans*; ZTRI_1.1705, an orthologue of *α*-agglutinin genes from *Fusarium sp*. and *Claviceps purpurea*, implicated in cell-surface adhesion [72]; ZTRI_3.465 and ZTRI_4.24, two further genes with orthologues in the *C. albicans* ALS family of agglutinins which function in cell-surface and cell-cell adhesion, each annotated as both induced and repressed in different kinds of *C. albicans* biofilms[72]; and ZTRI_9.291, an orthologue of biofilm-induced alcohol dehyrogenase genes in *C. albicans*[72]. All primers used in this study are detailed in Table 2.

**Table 2.**
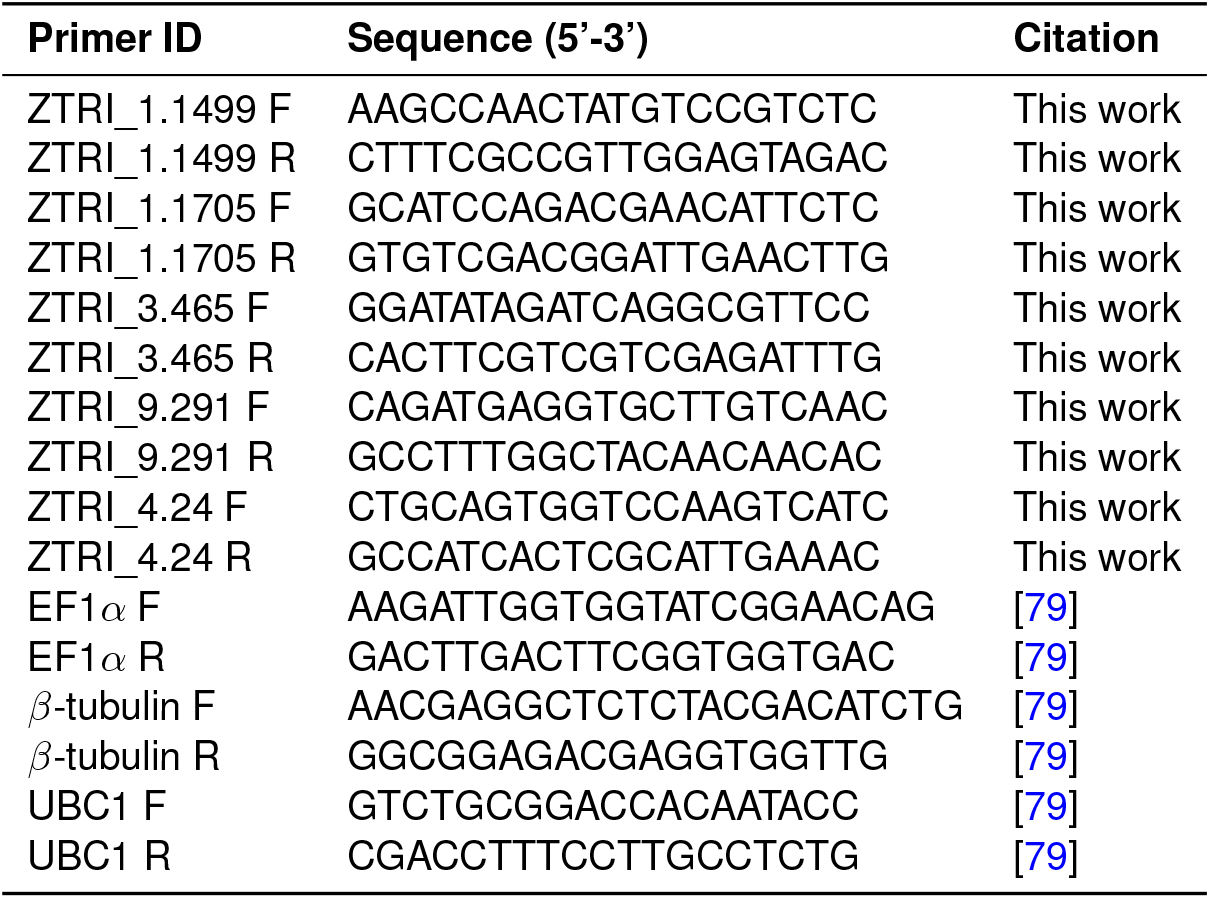
Primers used in this study.

qRT-PCR was performed in clear 384-well plates (Ambion, UK) using SYBR green in a 7300 Realtime PCR machine (Applied Biosystems). Relative expression of the genes of interest was normalised against the geometric mean expression of three reference genes (*EF1α, β-tubulin* and *UBC1*). Gene expression was calibrated using a standard curve of template concentration[85], produced using serial dilutions of pooled cDNA from all treatments. For each point on the standard curve, three technical replicates were used. All standard curves gave R^2^ values of at least 0.98. Gene expression was analysed according to the methods developed by Pfaffl *et al*.[86]. Three technical replicates, plus one no-RT control, were used per sample. No template controls were used for each primer pair, and dissociation curves were checked for evidence of mispriming or primer dimers. No-RT controls were used for all templates. The qRT-PCR experiment was carried out on two independent occasions.

## Author Contributions

TET carried out experiments and analysed data; GT carried out RNA extractions and RT-qPCR; CH prepared samples for SEM and oversaw this microscopy; HF conceived and oversaw the study, carried out experiments, analysed data and wrote the manuscript. All authors read and commented on the manuscript.

## Acknowledgments

We thank the University of Exeter’s A2I scheme and the BSPP for funding the work of Ms. Tegan Tyzack. We also thank the Technical Support Team associated with the University of Exeter’s Biosciences Mezzanine 2021-2022 for their tireless support of research: Ms Nicola Wood, Dr Nicola Senior, Dr Nasser Trissi and Mr James Smith.

## Supplementary Information

**Figure S1.**
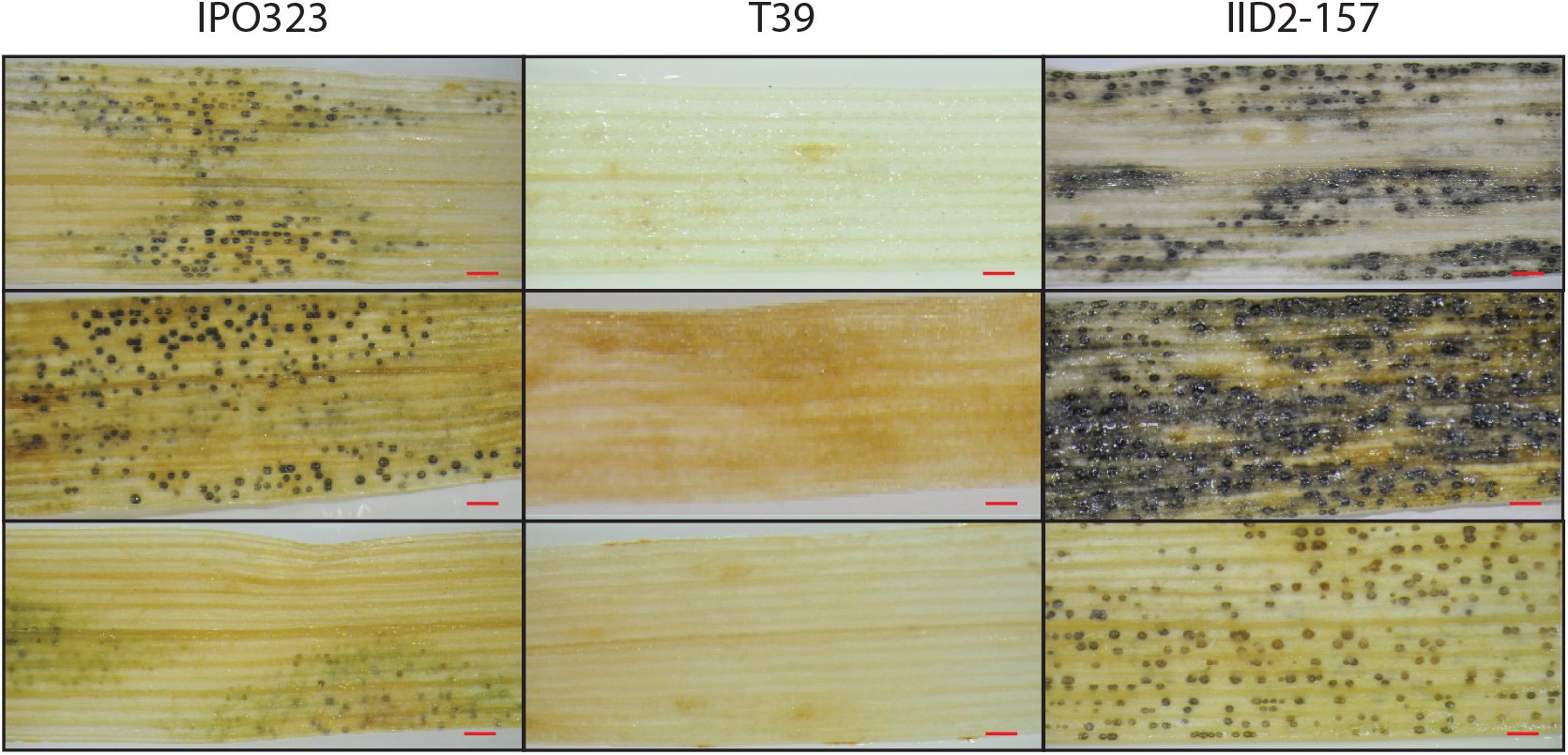
Figure S1: Ability of the three isolates, IPO323, T39 and IID2-157 to infect and complete their lifecycle on Triticum aestivam cultivar Galaxie (IPO323 and T39) or Triticum durum cultivar Volcani (IID2-157). IPO323 and IID2-157 are virulent on their respective hosts, while T39 is not able to produce pycnidia on Galaxie. Three images, each showing a 2.5 cm long field of view photographed with a macroscope at 2.5x magnification, are shown for each image. The images represent the variation seen across 2.5 cm leaf sections in each interaction. Scale bars represent 1 mm.

